# BMP suppresses WNT to integrate patterning of orthogonal body axes in adult planarians

**DOI:** 10.1101/2023.01.10.523528

**Authors:** Eleanor G. Clark, Christian P. Petersen

## Abstract

Adult regeneration restores patterning of orthogonal body axes after damage in a post-embryonic context. Planarians regenerate using distinct body-wide signals primarily regulating each axis dimension: anteroposterior Wnts, dorsoventral BMP, and mediolateral Wnt5 and Slit determinants. How regeneration can consistently form perpendicular tissue axes without symmetry-breaking embryonic events is unknown, and could either occur using fully independent, or alternatively, integrated signals defining each dimension. Here, we report that the planarian dorsoventral regulator *bmp4* suppresses the posterior determinant *wnt1* to pattern the anteroposterior axis. Double-FISH identified distinct anteroposterior domains within dorsal midline muscle that express either *bmp4* or *wnt1*. Homeostatic inhibition *bmp4* and *smad1* expanded the *wnt1* expression anteriorly, while elevation of BMP signaling through *nog1;nog2* RNAi reduced the *wnt1* expression domain. BMP signal perturbation broadly affected anteroposterior identity as measured by expression of posterior Wnt pathway factors, without affecting head regionalization. Therefore, dorsal BMP signals broadly limit posterior identity. Furthermore, *bmp4* RNAi caused medial expansion of the lateral determinant *wnt5* and reduced expression of the medial regulator *slit*. Double RNAi of *bmp4* and *wnt5* resulted in lateral ectopic eye phenotypes, suggesting *bmp4* acts upstream of *wnt5* to pattern the mediolateral axis. Therefore, *bmp4* acts at the top of a patterning hierarchy both to control dorsoventral information and also, through suppression of Wnt signals, to regulate anteroposterior and mediolateral identity. These results reveal that adult pattern formation involves integration of signals controlling individual orthogonal axes.

**Author Summary:** Systems that coordinate long-range communication across axes are likely critical for enabling tissue restoration in regenerative animals. While individual axis pathways have been identified, there is not yet an understanding of how signal integration allows repatterning across 3-dimensions. Here, we report an unanticipated linkage between anteroposterior, dorsoventral, and mediolateral systems in planarians through BMP signaling. We find that dorsally expressed BMP restricts posterior and lateral identity by suppressing distinct Wnt signals in adult planarians. These results demonstrate that orthogonal axis information is not fully independent and suggest a potentially ancient role of integrated axis patterning in generating stable 3-dimensional adult forms.

## Introduction

Most animal forms are organized along orthogonal body axes. Bilaterian body plans typically involve head-to-tail (anteroposterior, AP), back-to-belly (dorsoventral, DV), and midline-to-lateral (mediolateral, ML) dimensions produced with high fidelity. Signals defining each body axis can function largely independently, and many animals use canonical Wnt signaling to regulate the AP axis and BMP signaling to regulate the DV axis (1-3). Definitive body axes emerge through embryogenesis from initial asymmetries generated either by maternal cues such as oocyte polarity (4) or symmetry-breaking events such as sperm entry, cavitation, or gastrulation (3, 5). However, the ability of some species to undergo whole-body regeneration as adults suggests embryogenesis can be unnecessary to maintain and restore axis orthogonality. In planarians, acoels, and Cnidarians, patterning along individual tissue dimensions has been attributed to distinct spatial signaling pathways (6-13). Because such organisms can re-establish body forms for many successive generations asexually, they could in principle inherit AP and DV asymmetries from each axis dimension separately, leading to independence of orthogonal axis information. Alternatively, perpendicular axis systems might instead interact to enable coordinated growth. How separate patterning systems integrate across axes to generate three-dimensional pattern in adulthood is relatively unexplored.

The freshwater planarian *Schmidtea mediterranea* is a model for studying the principles of adult axis patterning due to its ability to regenerate nearly any surgically removed tissue and undergo perpetual homeostasis in the absence of injury. These abilities are supported by adult pluripotent stem cells termed neoblasts that differentiate into any of the approximately 150 cell types comprising the adult animal and assemble into functional and appropriately positioned and scaled tissues (14, 15). Neoblasts undergo regionalized specialization to form subsets of progenitors fated for tissues located at particular axial locations such as the eyes, pharynx, and dorsal-versus-ventral epidermal cells (16-18). Neoblasts also hone to particular regions through their migratory ability (19-22) in order to form organs at particular locations in the body (23-25). Therefore, spatial information is critical for controlling progenitor function in order to maintain and regenerate the planarian body plan.

Spatial organization in regeneration and homeostasis is provided by a specialized set of signaling factors termed position control genes (PCGs) that are expressed regionally from body-wall muscle (26). Following amputations that truncate the body axis, PCG expression domains shift to provide tissue identity information, allowing for restoration of missing body regions (27, 28). The planarian AP axis is controlled by canonical β-catenin signaling involving the use of posteriorly expressed Wnts and their signaling outputs, and anteriorly expressed Wnt-inhibitors (7, 9, 27-30). The Wnt ligand *wnt1* and secreted Wnt inhibitor *notum* are expressed at the posterior and anterior poles, respectively, where they organize tail and head patterning (7, 9, 27-29, 31). By contrast, the DV axis is established by BMP signaling. *bmp4* is expressed in a mediolateral gradient within dorsal muscle and acts to promote dorsal fates while repressing ventral fates, through feedback inhibition of a ventrally and laterally expressed *admp* homolog (6, 10, 32-35). Fates along the ML axis are determined through the reciprocal antagonism of medial *slit* and the laterally expressed non-canonical Wnt ligand *wnt5* (28, 36). *slit* inhibition results in a collapse of lateral tissues, such as eyes, onto the midline, while *wnt5* inhibition causes opposite defects in which ectopic tissue, such as eyes, form laterally. Inhibition of PCG factors results in mis-patterning phenotypes both in amputated animals regenerating a new blastema and also in uninjured animals using neoblasts to maintain their bodies through homeostasis. Therefore, canonical Wnt, BMP, and Slit/Wnt5 signals constitutively control axis identities across three spatial dimensions, but the independence versus interrelationship between these signals is not fully understood.

## Results

We sought to uncover possible relationships between key determinants of orthogonal body axes in planarians. We examined the expression of *bmp4* and observed that in addition to the prominent dorsal-versus-ventral expression pattern with highest expression on the dorsal midline (6, 10, 37), *bmp4* expression was stronger in the anterior versus posterior of the animal and reduced at the posterior tip (Fig. 1A). Furthermore, we noted that expression of *wnt1*, a master regulator of posterior identity, is selective to the dorsal and not ventral side of animals in the posterior tail approximately where *bmp4* expression is lower (9, 28) (Fig. 1A). *wnt1* is known to be co-expressed in a posterior subset of muscle cells specific to the dorsal midline marked by expression of *dd23400 (38)*. We utilized double fluorescence *in situ* hybridization (FISH) to examine the features of co-expression of dorsal posterior *wnt1*, dorsal *bmp4*, and dorsal midline *dd23400* (Fig. 1B). Similar to results reported previously, *dd23400* was co-expressed in 88.9% of *wnt1*+ cells (64/72 cells). Additionally, *dd23400* and *bmp4* were also co-expressed, as *bmp4* expression was detected in 55.4% of *dd23400*+ cells (169/305 cells) throughout the dorsal midline, though at apparently reduced numbers in the tail tip. By contrast, *bmp4* was only co-expressed in 12.1% of *wnt1*+ cells (4/33 cells) (Fig. 1B), with double-positive cells only present at the most anterior of the *wnt1* domain. Within the posterior *wnt1+* domain, *wnt1+* cells in the anterior of the domain co-expressed *bmp4* (3/18) while *wnt1* cells in the posterior of the domain were *bmp4*-negative (15/18). These results suggest a model in which *dd23400+* cells are partitioned into an anterior population co-expressing *bmp4* and a population in the far posterior co-expressing *wnt1*, with a limited overlap (Fig. 1C).

**Fig. 1.**
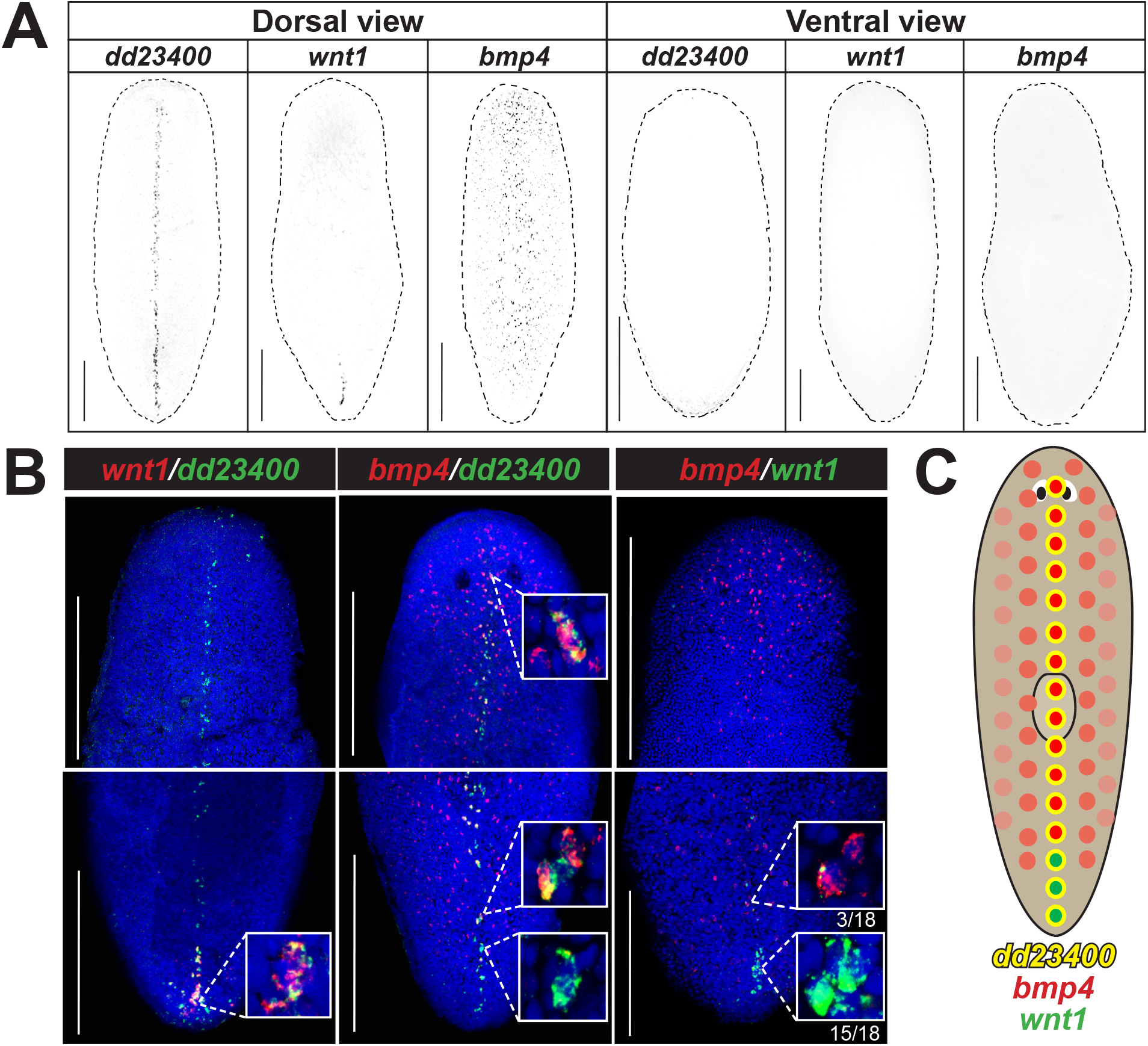
*bmp4* and *wnt1* co-express with *dd23400*+ dorsal midline cells in a regionally distinct manner. **(A)** Fluorescent *in situ* hybridization (FISH) detecting *dd23400* on the dorsal midline, *wnt1* on the dorsal posterior midline, and *bmp4* in a dorsal midline-centered gradient with reduced expression in the far posterior. Scale bars represent 150 μm with dorsal or ventral views indicated. **(B)** Double FISH detecting co-expression of *wnt1, dd23400*, and *bmp4*. 88.9% of *wnt1+* cells co-expressed *dd23400* and 55.4% of *dd23400+* cells co-expressed *bmp4*. Only 12.1% of *wnt1+* cells co-expressed *bmp4*. Therefore, along *dd23400+* dorsal midline muscle cells, *wnt1* and *bmp4* expression defines largely nonoverlapping AP domains within posterior. Within the *wnt1+* domain of the representative image, *wnt1+* cells in the anterior co-expressed *bmp4* (3/18) whereas cells in the posterior of the domain lacked *bmp4* co-expression (15/18). Scale bars represent 150 μm. **(C)** Schematic illustrating separation of *bmp4* and *wnt1* domains on the dorsal midline.

To test for possible functional relationships between *bmp4* and *wnt1*, we first used RNA interference (RNAi) to examine the consequences of *bmp4* inhibition. To circumvent the roles of *bmp4* in directing expression of the injury-induced *equinox* gene essential for blastema outgrowth (39), we examined axis relationships using homeostatic RNAi in the absence of injury to specifically reveal possible interactions between axis patterning factors. After 14 days of *bmp4* RNAi, the *wnt1* expression domain expanded dramatically anteriorly, but retained dorsally restricted specificity (Fig. 2A). Expanded expression of *wnt1* in *bmp4(RNAi)* animals was more sporadic and patchier compared to the normal *wnt1* domain in control animals. In addition, *wnt1* expression expanded laterally in *bmp4(RNAi)* animals. Together, these results indicate that *bmp4* controls the AP limit of *wnt1* expression.

**Fig. 2.**
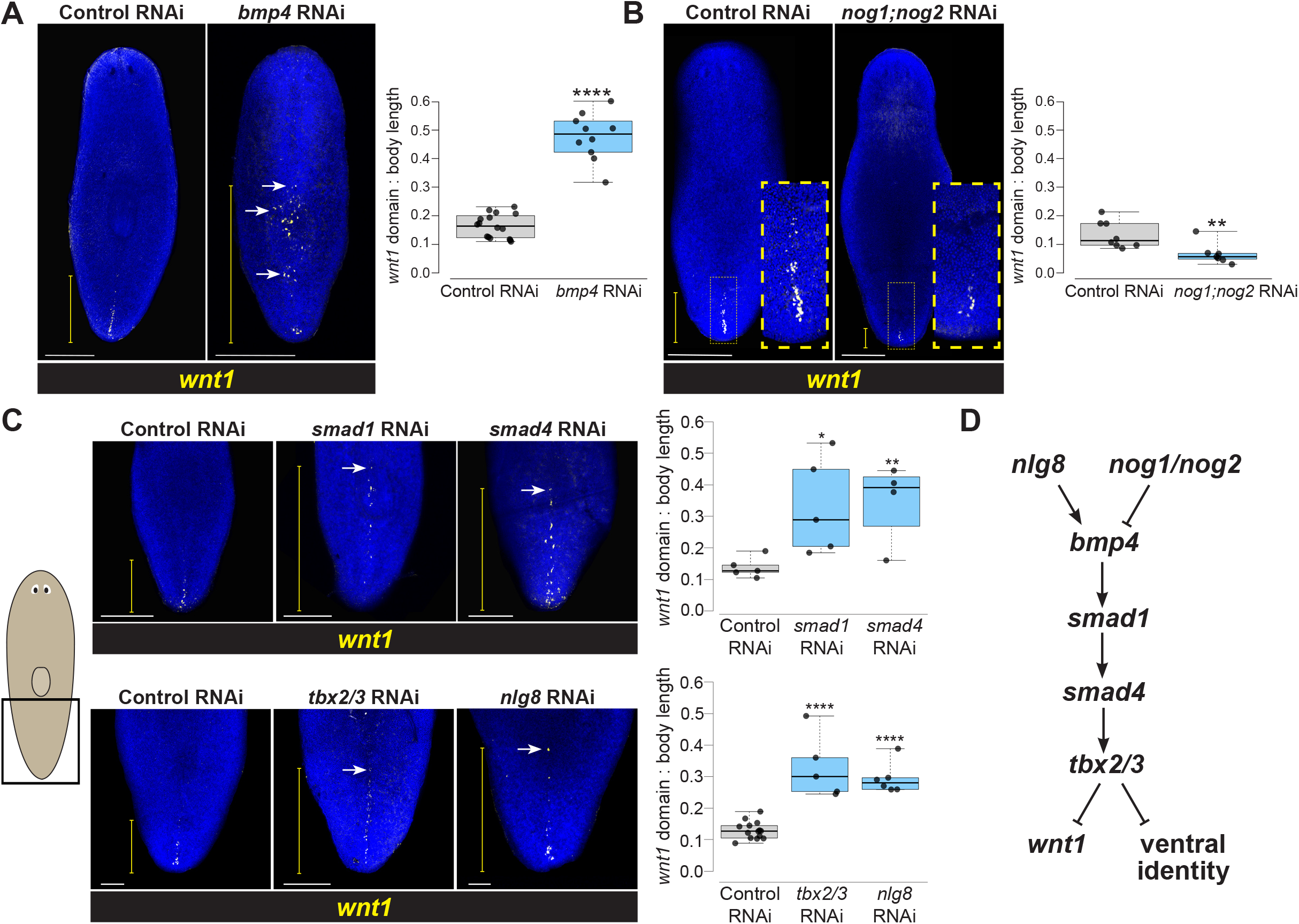
A BMP signaling pathway involved in DV identity restricts *wnt1* expression to the posterior. **(A)** FISH detecting *wnt1* expression in animals following 14 days of homeostatic control or *bmp4* RNAi. Inhibition of *bmp4* expanded *wnt1* expression toward the anterior (white arrow shows ectopic *wnt1+* cells). **(B)** *wnt1* expression domain reduced in *nog1;nog2(RNAi)* animals treated with dsRNA for 18 days of homeostasis (yellow boxes, magnified insets). **(C)** Top: FISH of *wnt1* following 14 days of control, *smad1*, or *smad4* RNAi. Bottom: FISH detecting *wnt1* after 18 days of control, *tbx2/3*, or *nlg8* RNAi. Inhibition of these BMP signaling components results in anterior expansion of *wnt1* expression. **(A-C)** White scale bars represent 150 μm, and yellow brackets indicate *wnt1* AP expression domain size. Box plots showing the distance from the posterior tip of the animal to the most anterior *wnt1* expression relative to body length after indicated treatments. *p<0.05, **p<0.01, ***p<0.001, ****p < 0.0001 by 2-tailed t-test; N ≥ 4 animals. Box plots show median values (middle bars) and first-to-third interquartile ranges (boxes); whiskers indicate 1.5× the interquartile ranges, and dots are data points from individual animals. **(D)** Pathway model for components acting in DV determination and control of *wnt1*.

To gain insights into whether BMP signaling acts permissively or instructively in regulation of *wnt1*, we inhibited Noggin homologs *nog1* and *nog2* known to negatively regulate *bmp4* in planarian DV determination (33). Homeostatic inhibition of *nog1* and *nog2* for 18 days decreased the *wnt1* expression domain (Fig. 2B), indicating that BMP signaling likely plays an instructive role in limiting posterior *wnt1* identity.

We next examined whether this role of *bmp4* in controlling a regulator of AP identity occurred via a canonical BMP pathway signaling through Smad1 and Smad4 effectors. Planarian *smad1* and *smad4* are known to mediate control dorsoventral identity along with *bmp4* (6, 10).

Following a 14-day inhibition, both *smad1(RNAi)* and *smad4(RNAi)* animals had significantly anterior expansion of *wnt1* expression, phenocopying the effects of *bmp4* RNAi on *wnt1* (Fig. 2C, top). Given these results, we next investigated potential regulation of *wnt1* by other factors known to act with BMP signaling to control dorsoventral identity. We examined the effects of inhibiting *tbx2/3*, a transcription factor hypothesized to act downstream of *bmp4* for control of DV identity in *Dugesia japonica* (40). Following 18 days of *tbx2/3* RNAi, *wnt1* expression was likewise significantly expanded toward the anterior (Fig. 2C, bottom). Similarly, we investigated *nlg8*, a noggin-like gene that facilitates BMP signal activation, is expressed dorsally, and whose inhibition phenocopies the ventralization phenotypes observed after *bmp4* RNAi (33).

Homeostatic RNAi of *nlg8* for 18 days caused expansion of the *wnt1* domain (Fig. 2C, bottom). Taken together, these experiments provide support that a *bmp4* signaling pathway closely linked to dorsoventral identity determination acts to restrict the expression domain of the posterior determinant *wnt1* (Fig. 2D).

To investigate whether the role of BMP signaling was limited to *wnt1* or more broadly affected posterior identity in general, we assessed expression of other posterior markers following 28 days of *bmp4* RNAi. Posterior markers *wnt11-2, wnt11-1, fzd4-1*, and *wntP-2* are expressed in successively broader posterior domains, have roles in tail and trunk patterning (7, 24, 30, 31, 41), and are expressed in a beta-catenin-dependent manner (42-44). Most known posterior markers such as these factors are expressed both dorsally and ventrally, unlike *wnt1*, but in regeneration their expression depends on *wnt1* (27, 28). *bmp4* inhibition expanded the domains of all four posterior markers in both their dorsal and ventral domains (Fig. 3A). Therefore, BMP signaling broadly limits posterior identity.

**Fig. 3.**
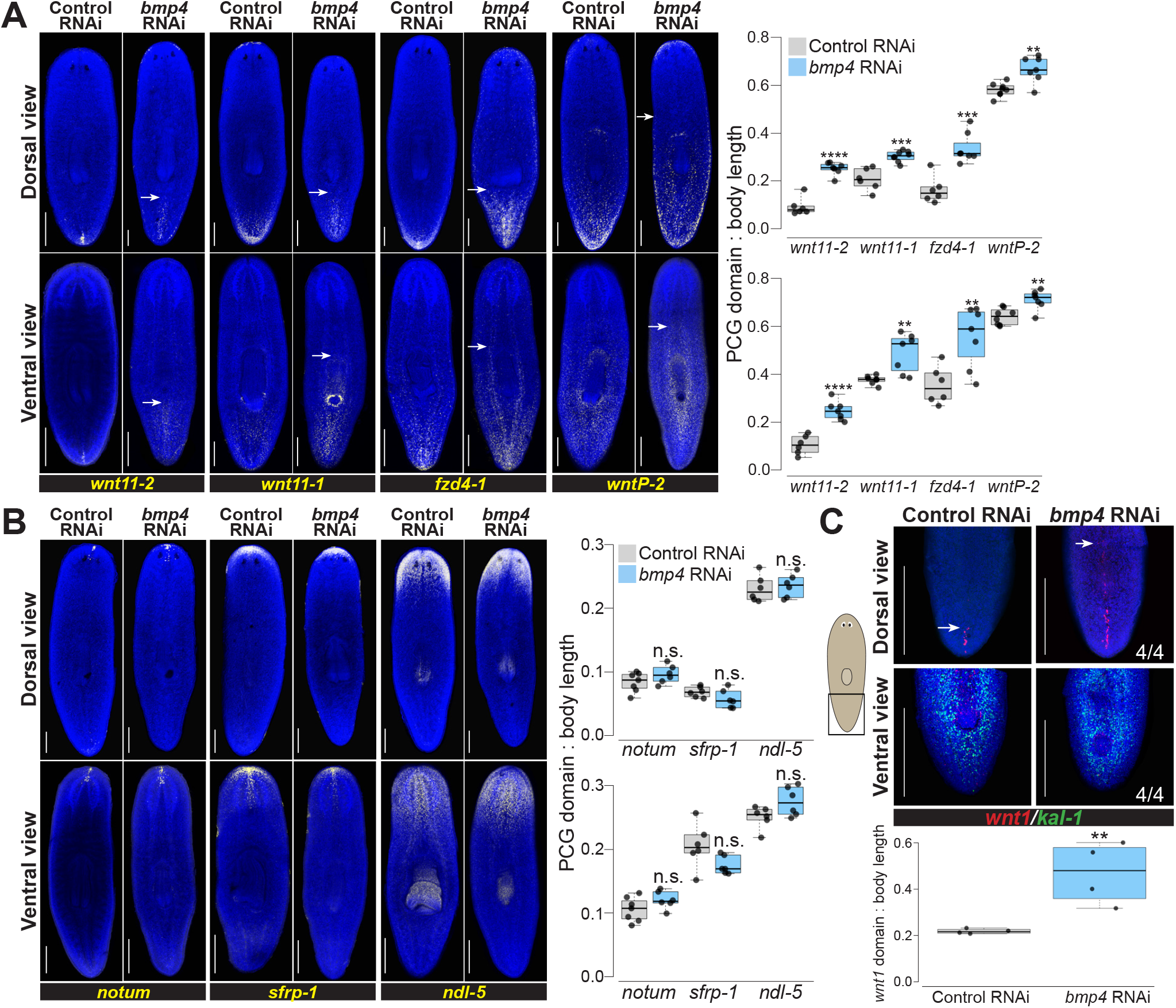
*bmp4* restricts posterior identity independent of DV control. **(A)** Animals were fixed after 28 days of homeostatic control or *bmp4* RNAi and stained by FISH as indicated for markers of the AP axis identity. *bmp4* RNAi caused an anterior expansion of the posterior markers *wnt11-2, wnt11-1, fzd4-1*, and *wntP-2* on both dorsal and ventral sides. White arrows indicate the expansion of posterior markers. **(B)** FISH stained animals following 28 days of control or *bmp4* RNAi. *bmp4* inhibition did not affect AP distribution of anterior markers *notum, sfrp-1*, or *ndl-5*. **(A-B)**, Graphs show the measurement of indicated marker domains normalized by total length of animal (N ≥ 6 animals). **(C)** FISH showing *wnt1* and ventral marker *kal-1* expression comparing 14 days of control and *bmp4(RNAi)*. Dorsal view (upper) shows that inhibition of *bmp4* expanded *wnt1* expression anteriorly (arrows indicate anterior-most *wnt1+* cell) at a time prior to any dorsal expression of the ventral marker *kal-1* expression (4/4 animals). Boxplot comparing *wnt1* expression normalized by body length between control and *bmp4(RNAi)* animals (N = 4 animals). **(A-C)** Box plots shows median values (middle bars) and first-to-third interquartile ranges (boxes); whiskers indicate 1.5× the interquartile ranges and dots are data points from individual animals. *p<0.05, **p<0.01, ***p < 0.001, ****p<0.0001, n.s. indicates p>0.05 by 2-tailed t-test. Scale bars, 300 μm (A-B) or 150 μm (C).

By contrast, markers of far anterior identity did not appear to become restricted in *bmp4* RNAi. We stained *bmp4(RNAi)* animals for *notum, sfrp-1*, and *ndl-5* in order to assess AP identity over a range of the anterior region (7, 9, 28, 29). Compared to control animals, there was no significant change in these expression patterns in the AP direction on either dorsal or ventral side of the animals (Fig. 3B). We note, however, that *bmp4(RNAi)* animals eventually form dorsal cephalic ganglia and an extra set of dorsal eyes (10). These transformations may impact anterior pattern expression to some degree, and we noted that the domain of *ndl-5* expression, present throughout the head of normal animals, appeared mediolaterally modified after homeostatic *bmp4* RNAi. However, our data suggest BMP’s role on the AP axis is primarily to regulate identity within the posterior.

Because *bmp4(RNAi)* animals undergo a progressive ventralization, we considered the possibility that ventral tissue identity might indirectly influence *wnt1* expression in these animals. To ascertain possible relationships between ventralization and posteriorization phenotypes, we examined *bmp4(RNAi)* animals at an early time in their phenotypic progression after 14 days of dsRNA feeding, then simultaneously assessed both phenotypes. These animals had expanded *wnt1* but not yet dorsal expression of the ventral epidermal marker *kal-1* (Fig. 3C). However, longer-term *bmp4* RNAi ultimately results in the dorsal expression of *kal-1* as the epidermis becomes ventralized during tissue turnover (16). Therefore, our results suggest that the anterior *wnt1* expansion after *bmp4* RNAi is unlikely a secondary consequence of tissue ventralization and instead could represent a separate use of BMP signaling for planarian AP axis patterning.

Given that *bmp4* and *wnt1* expression were enriched in separate anterior and posterior domains, we further examined whether these genes might undergo mutual negative regulation. To test this possibility, we inhibited *wnt1* homeostatically, followed by FISH to detect expression of *bmp4*. Following 14 days of *wnt1* RNAi, tails began to retract and become bulged, as reported previously (38). In these animals, the *bmp4* expression pattern was altered to be expressed more highly in the posterior of the animal but retained its dorsal specificity (Fig. S1). Therefore, inhibition of *wnt1* permitted *bmp4* expression in the dorsal posterior tip of the animal in a domain normally expressing *wnt1*. Together with the prior results, these experiments suggest a reciprocal antagonism between *wnt1* and *bmp4* to define each other’s expression boundaries and consequently pattern the posterior.

*wnt1* also undergoes dramatic expression dynamics early in regeneration. Wound sites express *wnt1* in muscle cells early after wounding, and optimal injury-induced *wnt1* expression depends on *bmp4* signals to induce expression of the novel secreted factor *equinox*, which in turn activates many injury-induced genes (39). In addition, regenerating tail fragments undergo an extensive remodeling of pre-existing territories so that regeneration restores the overall body proportionality without restoring absolute size. In regenerating tail fragments, *wnt1* expression undergoes an initial anterior expansion along the midline by 18 hours post-amputation, followed by eventual restriction and re-establishment of a new AP axis through rescaling over several days (28). We tested whether *bmp4* inhibition would affect these regeneration-dependent behaviors of *wnt1* expression along the posterior midline in amputated tail fragments. In these animals, *bmp4* RNAi resulted in an anterior expansion of the *wnt1* domain from the homeostatic knockdown prior to amputation and so was present in animals fixed immediately after amputation (Fig. S2). By 18 hours, control tail fragments underwent an anterior expansion of *wnt1* along the midline, while *bmp4(RNAi)* tail fragments retained an expanded *wnt1* domain. However, *bmp4(RNAi)* animals underwent apparently normal resetting of *wnt1* domains during the rescaling period by 96 hours, similar to control animals. By contrast, a prior study found that inhibition of the STRIPAK complex factor *mob4* led to anteriorly expanded *wnt1* but loss of regeneration-induced rescaling of *wnt1* territories (38). Therefore, although *bmp4* negatively regulates *wnt1* homeostatically, it is unlikely that the reduction to the *wnt1* expression domain through regenerative rescaling occurs through control of *bmp4* under normal conditions. Furthermore, it is likely that *bmp4* and *mob4* act separately to control *wnt1* expression.

In light of the unexpected role of BMP signals in AP axis patterning, we next sought to clarify how *bmp4* participates in ML axis regulation. *bmp4* RNAi causes expansion lateral tissue (*19*) and also failure to produce lateral tissue after lateral amputations. In addition, *bmp4* RNAi causes transverse regeneration to proceed with midline indentations (6, 10), likely because of *bmp4*’s role in dorsoventrally positioning the *notum+* anterior pole during head blastema outgrowth (45). Furthermore, *bmp4(RNAi)* homeostasis animals form an extra set of eyes medially, consistent with this factor having additional roles in ML axis formation (6, 10). However, the relationship between *bmp4* and other ML axis patterning factors *slit* and *wnt5* is not fully understood. We next examined *bmp4’s* role in homeostatically maintaining midline marker expression. Following 28 days of *bmp4* RNAi, expression of the midline determinant *slit* was reduced, particularly in the posterior of the animal (Fig. 4A). These results suggested that *slit* might function downstream of *bmp4* for controlling midline information. Furthermore, inhibition of *bmp4* reduced and disrupted the *dd23400* midline expression pattern and expanded its expression domain laterally (Fig. S3A), suggesting BMP controls midline identity broadly. Taken together, these data suggest that *bmp4* promotes medial identity and is important for establishing the boundaries of medial territories.

**Fig. 4.**
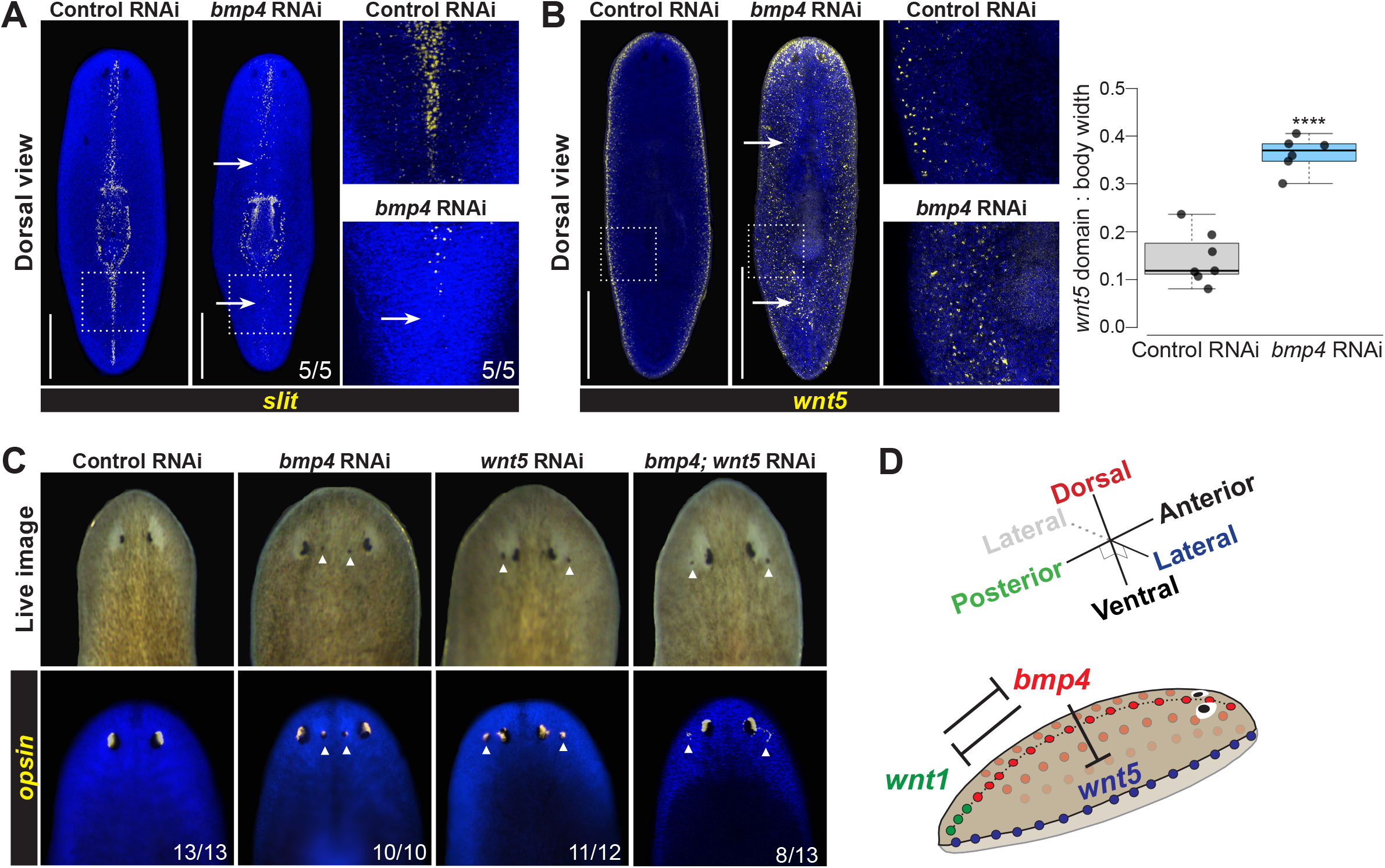
*bmp4* regulates ML patterning upstream of lateral *wnt5*. **(A-B)** FISH detecting *slit* or *wnt5* following 28 days of homeostatic control or *bmp4* RNAi. Right panels show enlargements of boxed regions. Scale bars represent 300 μm. **(A)** *bmp4* RNAi caused reduction of *slit* in the anterior and elimination in the posterior (arrows). **(B)** Inhibition of *bmp4* caused medial expansion of *wnt5* (arrows). Graph shows *wnt5* expression domain width (dorsal side) normalized to body width. ****p<0.0001 by unpaired 2-tailed t-test; N ≥ 6 animals. Box plots show median values (middle bars) and first to third interquartile ranges (boxes); whiskers indicate 1.5× the interquartile ranges and dots are data points from individual animals. **(C)** Eyes assessed by live imaging (top) or *opsin* FISH (bottom) after 21 days of homeostatic RNAi to inhibit *bmp* and/or *wnt5* and scored for lateral or medial ectopic eyes (arrows). For single-gene RNAi, dsRNA for the targeted gene was mixed with an equal amount of control dsRNA so that the total amounts of each targeted dsRNA delivered were equal between single-and double-RNAi conditions. 100% of *bmp(RNAi)* animals had ectopic medial eyes and 92% *wnt5(RNAi)* animals had ectopic lateral eyes. By contrast, 100% of *bmp4;wnt5(RNAi)* animals had at least two lateral ectopic eyes, and of these 38% also formed a single medial ectopic eye while 62% only formed ectopic lateral eyes (shown). Therefore, inhibition of *wnt5* masks the *bmp4(RNAi)* ML phenotype. **(D)** Model of *bmp4* establishing DV polarity, influencing AP regulation through *wnt1*, and controlling ML patterning through *wnt5*.

We next considered how *bmp4* might interact with lateral regulatory factor *wnt5*. We first used the lateral epidermal marker *laminB* to confirm prior results that *bmp4* inhibition generated ectopic lateral tissue (Fig. S3B) (32). We then examined a possible regulatory relationship between *bmp4* and the lateral determinant *wnt5*. Following 28 days of *bmp4* RNAi, expression of *wnt5* significantly expanded to occupy distant medial territories on the dorsal side (Fig. 4B), and more weakly expanded *wnt5* expression on the ventral side (Fig. S3C). To test whether *bmp4* and *wnt5* undergo reciprocal negative regulation, we inhibited *wnt5* for 21 days, then stained for *bmp4*. Inhibition of *wnt5* did not cause an apparent increase or decrease of *bmp4* expression under these conditions (Fig. S4). Together, these experiments demonstrate that *bmp4* antagonizes *wnt5* expression, promotes medial identity and suppresses lateral identity.

To determine whether *bmp4* functionally controls ML identity in part through regulation of *wnt5*, we conducted epistasis tests using eye placement as a readout. *bmp4* RNAi causes the formation of ectopic medial eyes, whereas *wnt5* RNAi produces an opposite defect of the formation of ectopic lateral eyes (6, 7, 10). To test for interactions between *bmp4* and *wnt5*, we homeostatically inhibited these genes individually or together for 21 days, examined animals visually, and stained them with an *opsin* riboprobe to label photoreceptor neurons (Fig. 4C).

Control animals had no ectopic eyes (13/13 animals), while 100% of *bmp4(RNAi)* animals (10/10 animals) had ectopic medial eyes, and 92% of *wnt5(RNAi)* animals (11/12 animals) had lateral eyes. In *bmp4;wnt5(RNAi)* animals, however, 100% of animals (13/13 animals) had at least two lateral ectopic eyes. Of these animals, 38% (5/13 animals) also had a single medial ectopic eye, while no animals displayed the *bmp4(RNAi)* phenotype of only medial ectopic eyes. Therefore, *wnt5* likely does not operate exclusively upstream of *bmp4*, because double-RNAi animals displayed the *wnt5(RNAi)* phenotype. Additionally, the two factors likely do not operate fully independently because *wnt5* co-inhibition reduced the penetrance of the *bmp4(RNAi)* phenotype (p=0.0027, 2-tailed Fisher’s exact test). Instead, taken together with the findings that *bmp4* RNAi causes expansion of *wnt5* expression, these results indicate *bmp4* can act upstream to limit *wnt5* in order to regulate ML identity.

## Discussion

Together, these experiments identify upstream roles for *bmp4* in patterning multiple body axes in planarians. BMP regulation not only establishes DV polarity but additionally influences both posterior identity through regulation of *wnt1* and also ML polarity through the suppression of *wnt5* and activation of *slit*. These results indicate that the signals governing perpendicular body axes (i.e., AP and DV axes) are not fully independent in adult pattern formation in planarians. We suggest that a regulatory logic in which the integration of information across axes may be important for robustness of patterning across long timescales in adulthood (Fig. 4D).

Our results clarify the relationships among the BMP, Wnt5, and Slit signals that participate in ML patterning in planarians. DV polarity is apparently normal in *wnt5(RNAi)* animals, suggesting this treatment does not strongly impact BMP-dependent patterning (Fig. S4)(28). Further, the *bmp4;wnt5(RNAi)* experiments presented here suggest *wnt5* acts downstream of *bmp4*, which is supported by *wnt5* expression expansion after *bmp4(RNAi)*. These results argue strongly against a hypothetical model in which *wnt5* acts exclusively upstream of *bmp4*. It has been previously demonstrated that BMP inhibits Wnt5 during ectoderm patterning in sea urchins (46) and BMP activity downregulates *wnt5a* to modulate convergence and extension in zebrafish development (47). Therefore, roles of BMP upstream of Wnt5 factors may be conserved. *slit* also likely acts downstream of *bmp4*, because *bmp4* inhibition reduced *slit* expression. These results collectively argue that *bmp4* acts at a high point in a hierarchy for ML patterning.

Our experiments also reveal interactions in which dorsal BMP restricts posteriorizing Wnt signals. These results are congruous with evidence of spatiotemporal coordination between the AP and DV axes in vertebrate development. In early mammalian development, a proximal BMP signal from trophectoderm activates Wnt3 asymmetrically in epiblast cells in order to establish the future AP axis (48). In early *Xenopus* development, BMP4 is required for the expression of posterior Wnt8 (49). The integration of BMP and Wnt signaling pathways occurs by Wnt8 preventing the conserved ability of GSK3 kinase to inhibit Smad1 activity through its phosphorylation and targeted proteasomal degradation, such that Wnt signals can positively enhance BMP signaling duration. This regulation contributes to a model in which AP positional information is specified by the duration of BMP signaling (50). Zebrafish development also involves coordination of AP and DV axis formation, with maternal Wnt signaling generating the organizer to repress *bmp*, and zygotic Wnt transcriptionally promoting the maintenance of BMP expression (51). Furthermore, BMP defines DV patterning in a temporal AP gradient from head to tail (52). By contrast, planarian Wnt/BMP integration involves signaling antagonism and operates at a transcriptional level and so likely occurs through a distinct mechanism. Uses for Wnt and BMP signals to pattern perpendicular body axes predate the bilaterians, with Cnidarians using Wnt signaling along the oral-aboral primary body axis and BMP to control the perpendicular directive axis (3, 12, 53, 54). In *Nematostella*, BMP and canonical Wnt signaling mutually antagonize to pattern endomesoderm (55). Our results suggest that interactions between BMP/Wnt signaling across axes may be a fundamental and ancient property enabling the integration of body axis information in three dimensions.

## Materials and Methods

### Experimental Model

Asexual *Schmidtea mediteraanea* animals (CIW4 strain) were kept in 1x Montjuic salts at 18-20°C. Animals were fed pureed calf liver. Animals were starved at least 7 days before the start of experiments.

### Fluorescent in situ hybridization (FISH)

The FISH protocol used is based off previously published work (38). Animals were killed in 7.5% NAC in PBS, fixed in 4% formaldehyde, and stored in methanol. They were rehydrated with methanol:PBSTx, bleached in 6% hydrogen peroxide in PBS, permeabilized with proteinase K, then prehybridized at 56°C. Hybridization occurred with digoxigenin or fluorescein labeled riboprobes at a 1:1000 concentration, which were synthesized using T7 RNA binding sites for antisense transcription. Animals were washed in a SSC concentration series at 56°C. Anti-digoxigenin-POD or anti-fluorescein-POD antibodies were in a solution of TNTx/ horse serum/ Western blocking reagent at a concentration of 1:2000. Tyramide in TNTx was utilized to develop and amplify the antibody signal. For double FISH, the enzymatic activity of tyramide reactions was inhibited by sodium azide. Nuclei were stained using 1:1000 Hoescht in TNTx.

### RNA interference (RNAi)

RNAi treatments were performed by dsRNA feeding with 80% liver and 5% food dye. dsRNA was synthesized as previously described (29). Animals were fed RNAi food every 2-3 days for the length of the experiment. For double RNAi, control dsRNA was mixed in with the single experimental dsRNA to ensure the same amount of overall dsRNA in feedings between double and single experimental conditions (Fig. 4C). For RNAi treatment without injury, animals were fixed 5 days after the last feeding. For a regeneration time course following injury, animals were cut 2 days after the last feeding and fixed at the indicated time.

### Image Acquisition

Live animals were imaged with a Leica M210F dissecting microscope with a Leica DFC295 camera (Fig. 4C). Stained animals were imaged with a Leica DMI 8 confocal microscope (Fig. 1-4, S1-S4). FISH images are maximum projections from a z-stack. Adjustments to brightness and contrast were made using Adobe Photoshop or ImageJ.

### Primer Design

Primers for dsRNA and riboprobes are listed in Table S1.

### Quantification and Statistical Analysis

Body length and PCG measurements were taken using LAS X or FIJI on z-stack maximum projections of FISH images. For body length, animals were measured from their most anterior to most posterior tips. For PCG length, animals were measured from the tip of the anterior (Fig. 3B), tip of the posterior (Fig. 2, 3A, 3C, S2), or lateral edge (Fig. 4B) to end of the fluorescence stain. Cell counting was manually performed using maximum projections in LAS X (Fig. 1, S3). Statistical analysis was conducted using Microsoft Excel. Plots were generated in BoxPlotR.

## Acknowledgements

We thank members of the Petersen lab for critical comments and Dr. Erik Schad for reagents and concepts. This work was supported by National Institutes of Health grant NIGMS R01GM129339 (to C.P.P.), National Institutes of Health grant NIGMS R01GM130835 (to C.P.P.), Simons/SFARI (597491-RWC) pilot project grant (to C.P.P.). E.G.C was supported in part by the Northwestern University Graduate School Cluster in Biotechnology, Systems, and Synthetic Biology, which is affiliated with the Biotechnology Training Program, and Northwestern University biotechnology training grant.

## Data and materials availability

All data are available in the main text or the supplementary materials.

## Supplementary Figure Legends

**Fig. S1.**
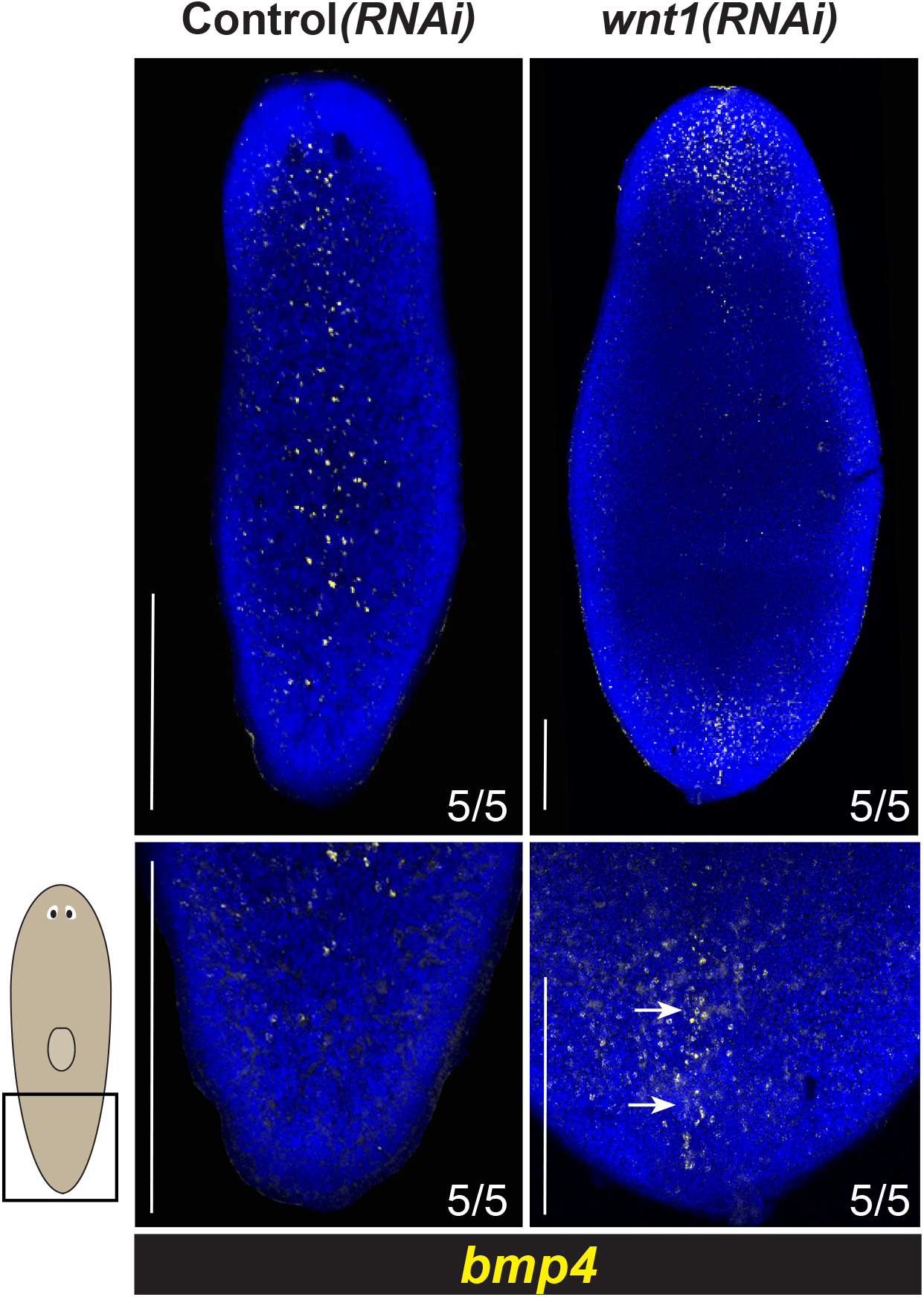
*wnt1* inhibits *bmp4* expression in the posterior. FISH staining for *bmp4* expression after 14 days of control or *wnt1* RNAi. Inhibition of *wnt1* results in elevated expression of *bmp4* on the posterior midline of the animal (arrows). Bottom panels show enlargements. Scale bars are 150 μm. N = 5 animals.

**Fig. S2.**
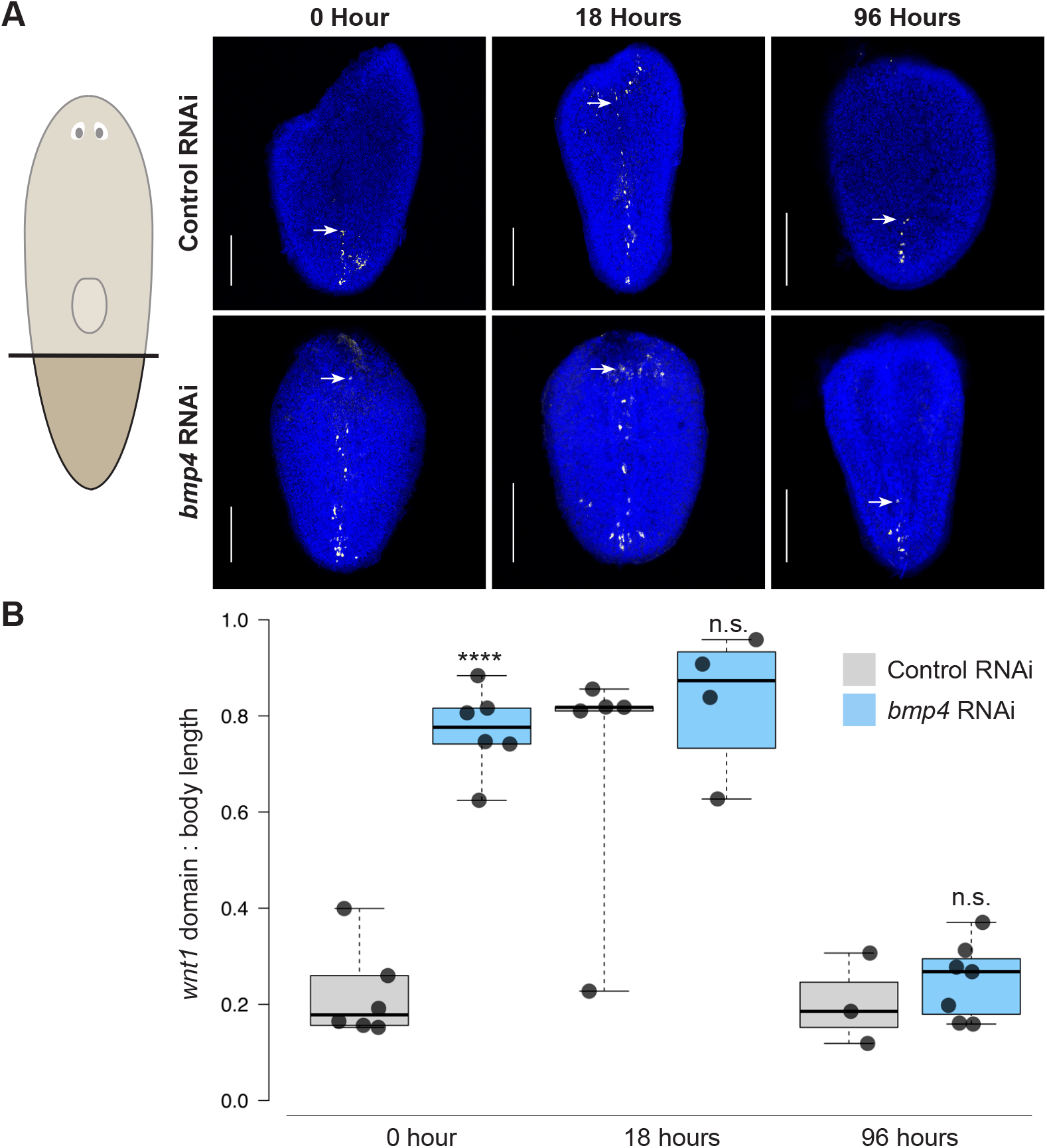
Reduction of *wnt1* expression domain through AP regenerative rescaling is not influenced by *bmp4* inhibition. **(A)** FISH to detect *wnt1* expression in regenerating tail fragments at 0, 18, and 96 hours after amputations conducted after 14 days of either control or *bmp4* RNAi. Arrows indicate anterior-most *wnt1+* cell detected along the dorsal midline for each timepoint and condition. Top panels show control animals undergoing early expansion of the dorsal midline *wnt1* domain by 18 hours of regeneration followed by rescaling to reduce the domain to the tip of the animal by 96 hours. Bottom panel shows *wnt1* expression dynamics in *bmp4* RNAi, in which midline *wnt1* expression was anterior expanded at the time of injury (0 hours), remained expanded at 18 hours of regeneration, and then restricted posteriorly by 96 hours, similar to control RNAi conditions. Therefore, BMP pathway modulation is unlikely to be responsible for the normal restriction of *wnt1* by 96 hours in regenerating tail fragments. Scale bars represent 150 μm. **(B)** Graph showing the quantification of the length of the *wnt1* domain relative to length of tail fragment. ****p<0.0001 by 2-tailed t-test and n.s. indicates p>0.05; N ≥ 3 animals. Box plots shows median values (middle bars) and first to third interquartile ranges (boxes); whiskers indicate 1.5× the interquartile ranges and dots are data points from individual animals.

**Fig. S3.**
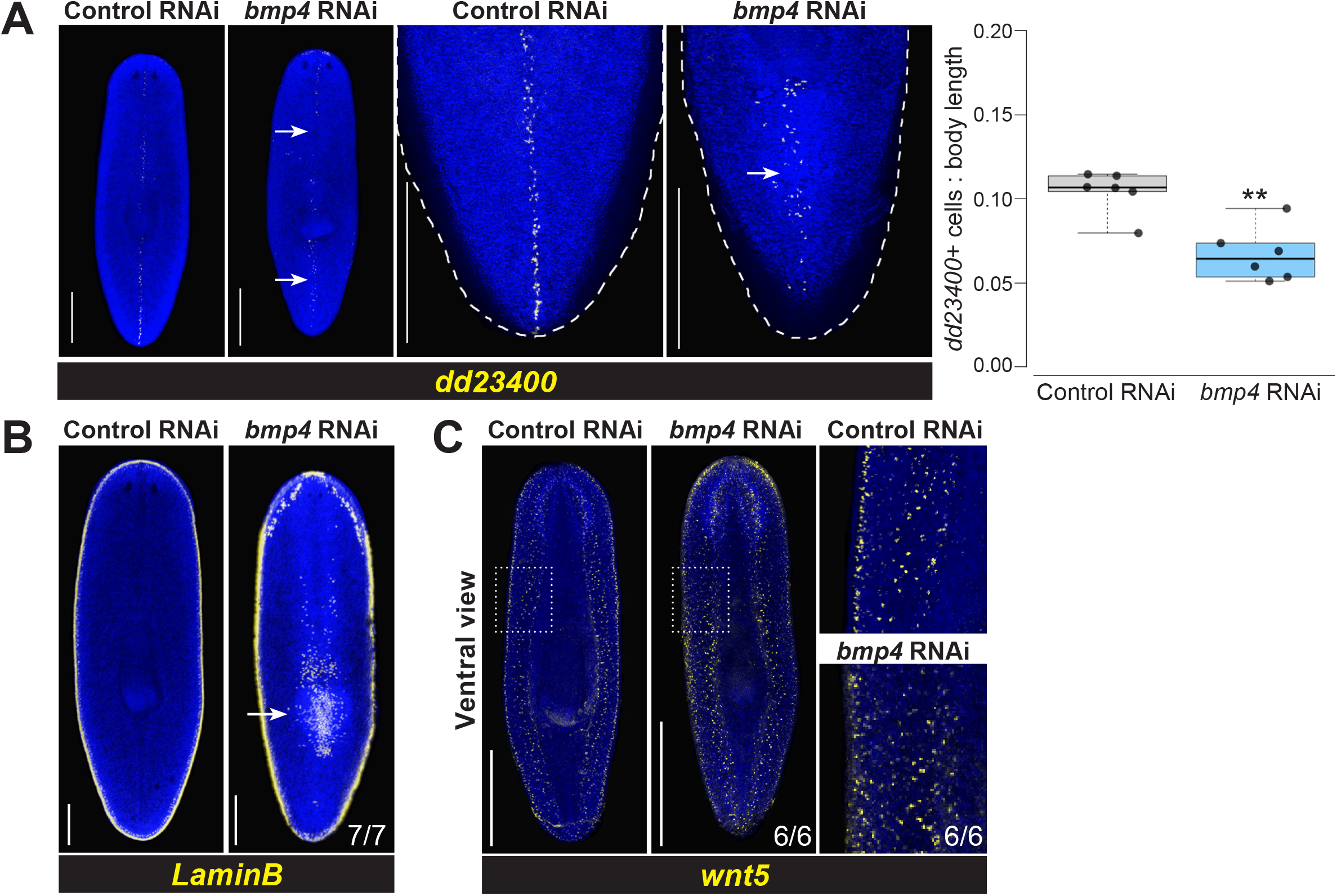
*bmp4* promotes midline identity and suppresses lateral identity. **(A-C)** FISH for *dd23400, LaminB*, and *wnt5* following 28 days of control or *bmp4* RNAi. Scale bars represent 300 μm. **(A)** Inhibition of *bmp4* reduces *dd23400* expression (arrows), particularly in the posterior. Right: Quantification of number of *dd23400+* cells normalized to animal body length. **p<0.01 by unpaired 2-tailed t-test; N ≥ 6 animals. Box plots show median values (middle bars) and first to third interquartile ranges (boxes); whiskers indicate 1.5× the interquartile ranges and dots are data points from individual animals. **(B)** *bmp4* RNAi causes ectopic medial expression of lateral marker *laminB* expression on the posterior midline (arrows). **(C)** Knockdown of *bmp4* appears to elevate *wnt5* expression less dramatically on the ventral side versus dorsal side. Right panels show enlargements of boxed regions.

**Fig. S4.**
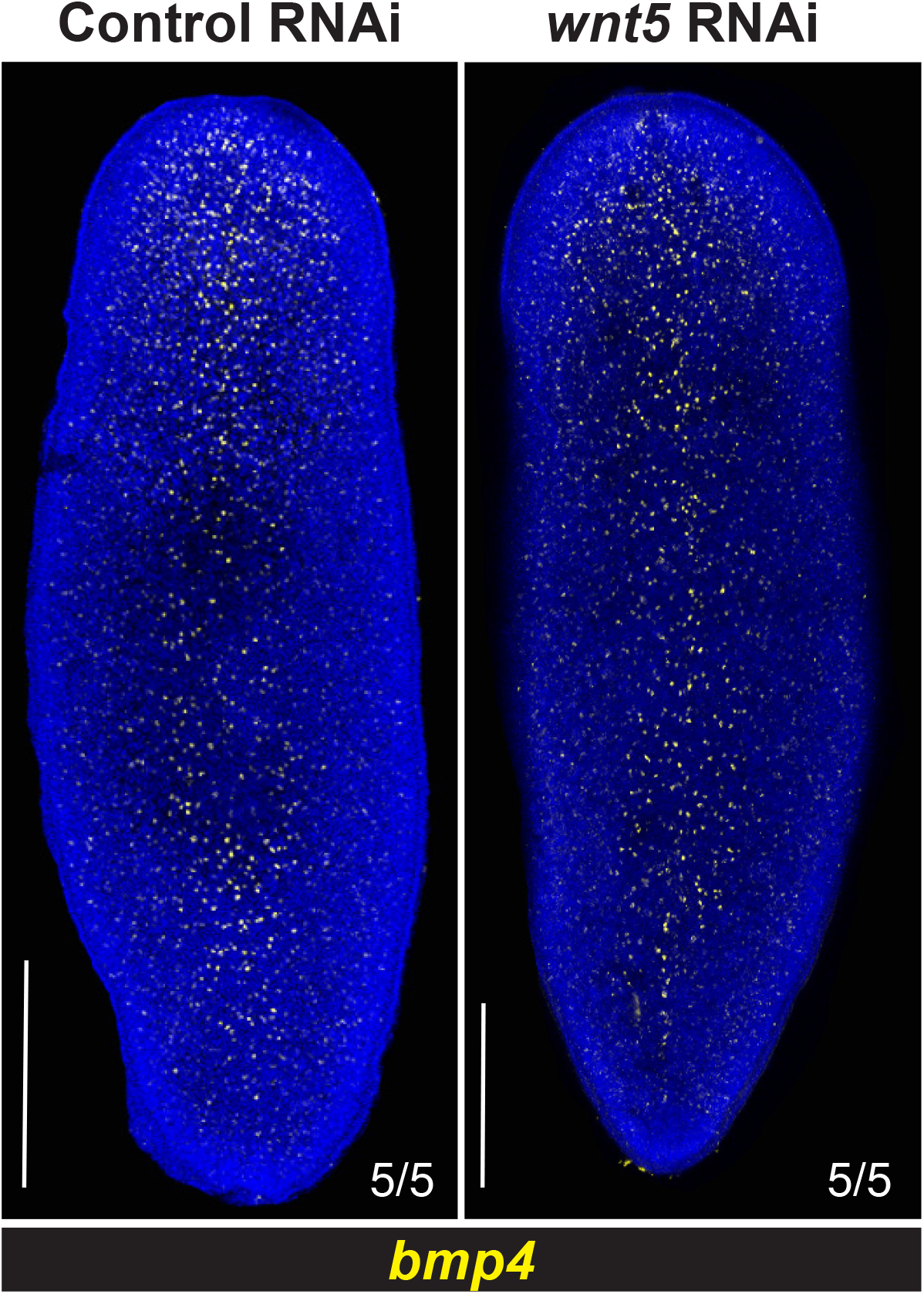
*wnt5* inhibition does not detectably alter *bmp4* expression. FISH staining for *bmp4* after 21 days of control or *wnt5* RNAi under homeostatic conditions. Inhibition of *wnt5* does not cause detectable increases or decreases in *bmp4* expression or distribution. Scale bars are 150 μm. N = 5 animals.

**Table S1.**
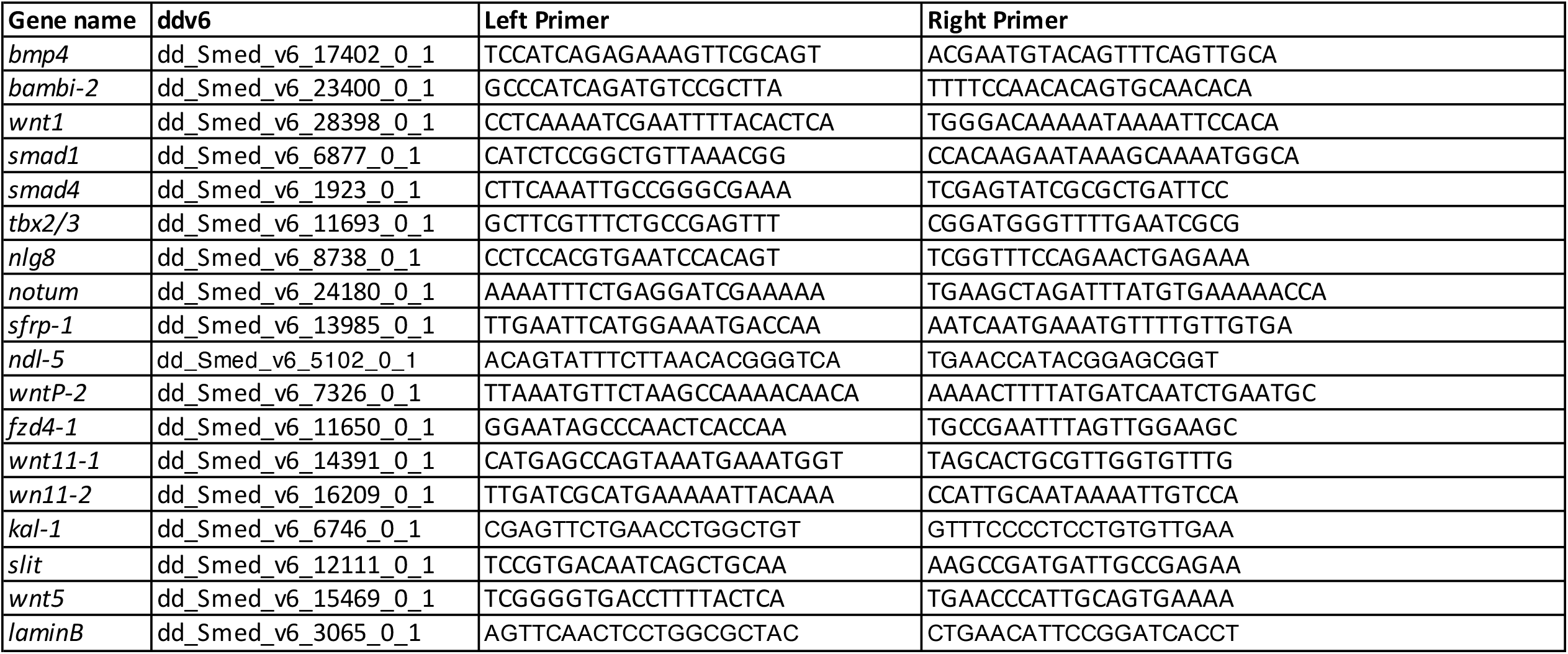
Primer Sequences for dsRNA and riboprobes.

